# Developing an automated iterative near-term forecasting system for an ecological study

**DOI:** 10.1101/268623

**Authors:** Ethan P. White, Glenda M. Yenni, Shawn D. Taylor, Erica M. Christensen, Ellen K. Bledsoe, Juniper L. Simonis, S. K. Morgan Ernest

## Abstract

1. Most forecasts for the future state of ecological systems are conducted once and never updated or assessed. As a result, many available ecological forecasts are not based on the most up-to-date data, and the scientific progress of ecological forecasting models is slowed by a lack of feedback on how well the forecasts perform.
2. Iterative near-term ecological forecasting involves repeated daily to annual scale forecasts of an ecological system as new data becomes available and regular assessment of the resulting forecasts. We demonstrate how automated iterative near-term forecasting systems for ecology can be constructed by building one to conduct monthly forecasts of rodent abundances at the Portal Project, a long-term study with over 40 years of monthly data. This system automates most aspects of the six stages of converting raw data into new forecasts: data collection, data sharing, data manipulation, modeling and forecasting, archiving, and presentation of the forecasts.
3. The forecasting system uses R code for working with data, fitting models, making forecasts, and archiving and presenting these forecasts. The resulting pipeline is automated using continuous integration (a software development tool) to run the entire pipeline once a week. The cyberinfrastructure is designed for long-term maintainability and to allow the easy addition of new models. Constructing this forecasting system required a team with expertise ranging from field site experience to software development.
4. Automated near-term iterative forecasting systems will allow the science of ecological forecasting to advance more rapidly and provide the most up-to-date forecasts possible for conservation and management. These forecasting systems will also accelerate basic science by allowing new models of natural systems to be quickly implemented and compared to existing models. Using existing technology, and teams with diverse skill sets, it is possible for ecologists to build automated forecasting systems and use them to advance our understanding of natural systems.

## Introduction

Forecasting the future state of ecological systems is important for management, conservation, and evaluation of how well models capture the processes governing ecological systems (Clark et al., 2001; Tallis & Kareiva, 2006; Díaz et al., 2015; Dietze, 2017). In 2001, Clark et al. (2001) called for a more central role of forecasting in ecology. Since then, an increasing number of ecological forecasts are being published that focus on societally important questions from daily to decadal time scales (Dietze et al., 2018). At daily scales, ecological forecasts predict the occurrence of environmental issues like toxic algal blooms (Stumpf et al., 2009) and pollen (Prank et al., 2013). At monthly scales, forecasts are used to predict the stocks of fisheries (NOAA, 2016) and the probability of coral bleaching events (Liu et al., 2018). At decadal time scales, ecological forecasts are used to predict how biodiversity will change as it responds to anthropogenic influences (Harris et al., 2018). These forecasting examples highlight the important role that ecological forecasts play in recasting ecological knowledge in societally relevant ways and also improve our understanding of ecological systems by testing the ability of our models to predict how systems will change in the future (Dietze et al., 2018; Harris et al., 2018).

While some of the examples given above (e.g., fisheries stock estimates) are regularly repeated, most ecological forecasts are made once, published, and never assessed or updated (Dietze et al., 2018). This lack of both regular assessment and active updating has limited the progress of ecological forecasting and hindered our ability to make useful and reliable predictions. The lack of active assessment results in limited information on how much confidence to place in forecasts and makes it difficult to determine on which forecasting methods to build. Without regular updates, forecasts lack the most current data, and the longer a forecast remains out of date, the less accurate it becomes (Petchey et al., 2015; Dietze et al., 2018). More regular updating and assessment will advance ecological forecasting as a field by accelerating the identification of the best models for individual forecasts and improving our understanding of how to best design forecasting approaches for ecology in general. This approach has helped accelerate forecasting ability in other fields such as meteorology (Kalnay, 2003; McGill, 2012; Bauer et al., 2015). For ecological forecasting to mature as a field, we need to change how we produce and interact with forecasts, creating a more dynamic interplay between model development, prediction generation, and incorporation of new data and information (Dietze et al., 2018).

With the goal of making ecological forecasting more dynamic and responsive, Dietze et al. (2018) recently called for an increase in iterative near-term forecasting. Iterative near-term forecasting is defined as making predictions for the near future and repeatedly updating those predictions through a cycle of evaluation, integration of new data, and generation of new forecasts. Because forecasts are made ‘near-term’—daily to annual time scales instead of multi-decadal—predictions can be assessed more quickly and frequently, leading to more rapid model improvements (Tredennick et al., 2016; Dietze et al., 2018). Since forecasts are made repeatedly through time, new data can be continuously integrated with each iteration (Dietze et al., 2018). By quickly identifying how models are failing, facilitating rapid testing of improved models, and incorporating the most up-to-date data available, iterative near-term forecasting has the potential to promote rapid improvement in the state of ecological forecasting. In addition to yielding improved information for guiding policy and management (Clark et al., 2001; Luo et al., 2011; Petchey et al., 2015), this iterative approach will help improve our basic understanding of ecological systems (Dietze et al., 2018). For example, alternative mechanistic models can be compared to determine which model provides the best forecasts, thus providing insights into the importance of different ecological processes (Dietze et al., 2018). Iterative near-term forecasting provides the more dynamic interplay between models, predictions, and data that has been identified as necessary for improving ecological forecasting and our understanding of ecological systems more broadly.

Because iterative near-term forecasting requires a dynamic integration of models, predictions, and data, Dietze et al. (2018) highlight approaches to data management, model construction and evaluation, and cyberinfrastructure that are necessary to effectively implement this type of forecasting (Box 1). Data needs to be released quickly under open licenses (Vargas et al., 2017; Dietze et al., 2018) and structured so that it can be used easily by a variety of researchers and in multiple modeling approaches (Borer et al., 2009; Strasser et al., 2011). Models need to be able to deal with uncertainty, in both the predictors and the predictions, to properly convey uncertainty in the resulting forecasts (Diniz-Filho et al., 2009). Multiple models should be developed, both to assess which models are performing best (Dietze et al., 2018) and to facilitate combining models to form ensemble predictions which tend to perform better than single models (Araujo & New, 2007; Diniz-Filho et al., 2009; Dormann et al., 2018). Ensuring that data and models are regularly updated and new forecasts are made requires cyberinfrastructure to automate data processing, model fitting, prediction, model evaluation, forecast visualization, and archiving. In combination, these approaches should allow forecasts to be easily rerun and evaluated as new data becomes available (Box 1; Dietze et al., 2018).

While iterative near-term forecasting is an important next step in the evolution of ecological forecasting, the requirements outlined by Dietze et al. (Box 1) are not trivial to implement (e.g., making quality data available in near real-time and automatically rerunning forecasts in reproducible ways), and few of their recommendations are in widespread use in ecology today (e.g., Wilson et al., 2014; Stodden & Miguez, 2014; Yenni et al., 2018). We explored what it would entail to operationalize Dietze et al’s recommendations by constructing our own iterative near-term forecasting pipeline for an on-going, long-term ecological study that collects high-frequency data on desert rodent abundances (Brown, 1998; Ernest et al., 2008). We constructed an automated forecasting pipeline with the goal of being able to forecast rodent abundances and evaluate our predictions on a monthly basis. In this paper, we discuss our approach for creating this iterative near-term forecasting pipeline, the challenges we encountered, the tools we used, and the lessons we learned so that others can create their own iterative forecasting systems. For those interested in implementing iterative forecasting, either on their own or as part of a team, this paper will provide a roadmap for how to build such a system and what skills will be helpful to do so. For readers looking for an introduction to automation and continous integration in an ecological context, we recommend our paper on data management for continuously collected data, which includes a tutorial on how to set up some of the aspects of automation described in this paper (Yenni et al., 2018).

## System Background

Iterative forecasting is most effective with frequently collected data, since it provides more opportunities for updating model results and assessing (and potentially improving) model performance (Box 1; Dietze et al., 2018). The Portal Project is a long-term ecological study situated in the Chihuahuan Desert (2 km north and 6.5 km east of Portal, Arizona, US). Researchers have been continuously collecting data at the site since 1977, including data on the abundance of rodent and plant species (monthly and twice yearly, respectively) and climatic factors such as air temperature and precipitation (daily) (Brown, 1998; Ernest et al., 2009, 2016, 2018). The site consists of 24 50m x 50m experimental plots. Each plot contains 49 permanently marked trapping stations laid out in a 7 x 7 grid, and all plots are trapped with Sherman live traps for one night each month. For all rodents caught during a trapping session, information on species identity, size, and reproductive condition is collected, and new individuals are given identification tags. This information on rodent populations is high-frequency, uses consistent trapping methodology, and has an extended time-series (475 monthly samples and counting), making this study an ideal case for near-term iterative forecasting. Forecasting of rodent population dynamics in the southwest (and more broadly) is important because of their link to zoonotic diseases such as hantavirus and plague (Parmenter et al., 1993; Gage & Kosoy, 2005; Springer et al., 2016). In addition, this forecasting system is being used to improve population dynamic modeling for this community and to explore the utility of incorporating experimental data into ecological forecasting models.

## Implementing an automated iterative forecasting system

Implementation of iterative forecasting requires the regular updating of models with new raw data as it becomes available and the presentation of those forecasts in usable forms; in our case, this occurs monthly. Updating models in an efficient and maintainable way relies on developing an automated pipeline to handle the six stages of converting raw data into new forecasts: data collection, data sharing, data manipulation, modeling and forecasting, archiving, and presention of the forecasts (Figure 1a). To implement the pipeline outlined in Figure 1a, we used a “continuous analysis” framework (*sensu* Beaulieu-Jones & Greene, 2017) that automatically processes the most up-to-date data, updates the models, makes new forecasts, archives the forecasts, and updates a website with analysis of current and previous forecasts. In this section we describe our approach to streamlining and automating the multiple components of the forecasting pipeline and the tools and infrastructure we employed to execute each component.

**Figure 1:**
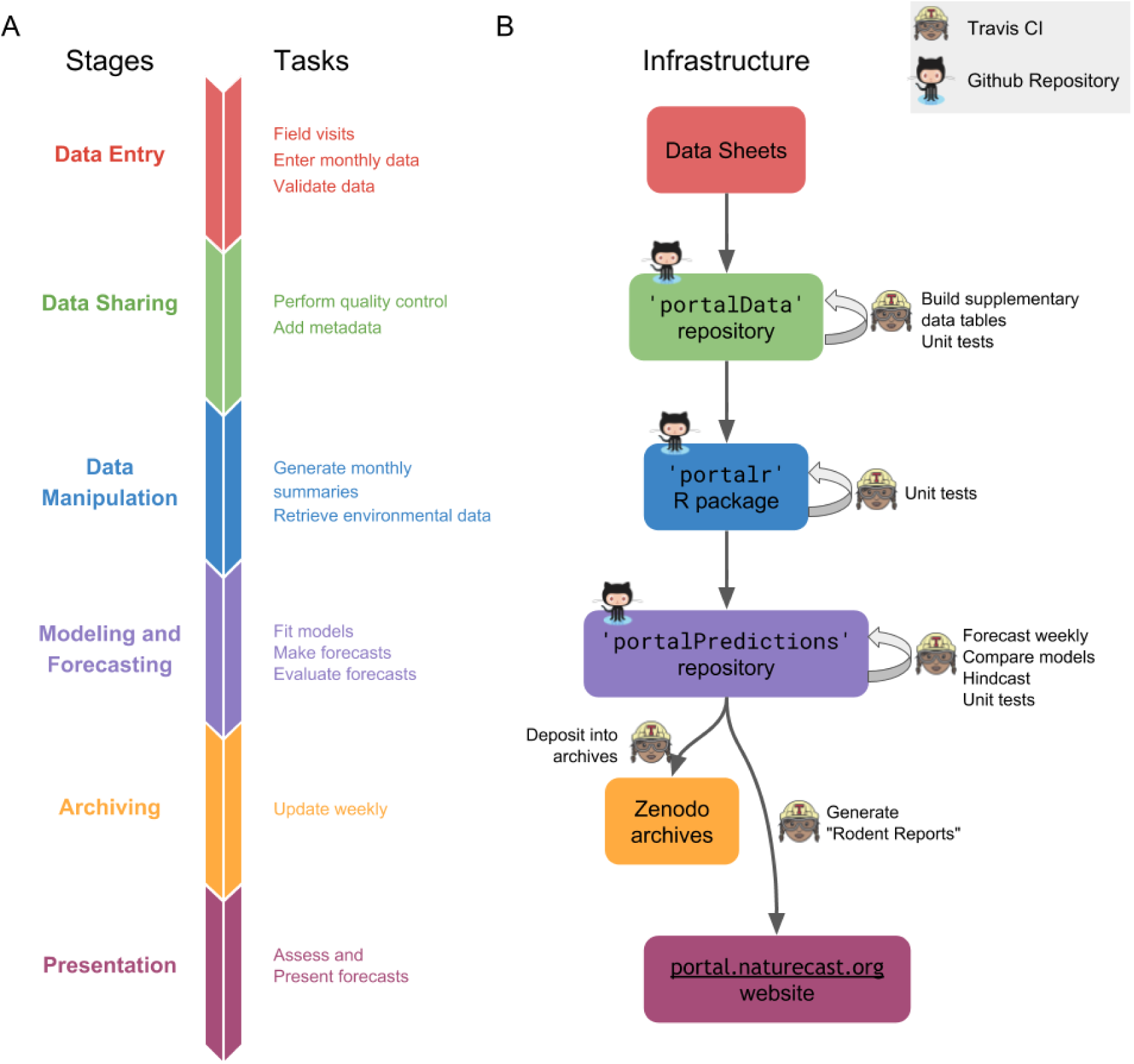
A) Stages of the forecasting pipeline. To go from raw data to forecast presentation involves a number of stages, each of which requires unique tasks, tools and infrastructure. The stages are interdependent, with outputs from one stage forming the inputs for the subsequent stage. Tasks in all stages are run using code written in R. B) Continuous integration system. Each box denotes the core infrastructure used for each stage of the forecasting pipeline. Continuous integration (denoted by the Travis icon, a woman wearing safety glasses and hardhat) triggers the code involved in events that link the stages of the pipeline, such as using the output from the forecasting stage (purple box) to create an updated website (rose box). Travis also runs tasks within a stage, such as testing code and adding weather data (icons on arrows originating and ending on the same box). The code for driving different stages of this pipeline is stored on GitHub (denoted by the GitHub icon, an “octocat”).

## Continuous Analysis Framework

A core aspect of iterative near-term forecasting is the regular rerunning of the forecasting pipeline. We employed “continuous analysis” (*sensu* Beaulieu-Jones & Greene, 2017) to drive the automation of both the full pipeline and a number of its individual components. Continuous analysis uses a set of tools originally designed for software development called “continuous integration” (CI). CI combines computing environments for running code with monitoring systems to identify changes in data or code. Essentially, CI is a computer helper who watches the pipeline and, when it sees a change in the code or data, runs all the computer scripts needed to ensure that the forecasting pipeline runs from beginning to end. This is useful for iterative near-term forecasting because it does not rely on humans to create new forecasts whenever new models or data are added. These tools are common in the area of software development, where they are used to automate software testing and integrate work by multiple developers working on the same code base. However, these tools can be used for any computational task that needs to be regularly repeated or run after changes to code or data (Beaulieu-Jones & Greene, 2017). Our forecasting pipeline currently runs on a publicly available continuous integration service (Travis CI; https://travis-ci.org/) that is free for open source projects (up to a limited amount of computing time). This continuous integration integrates directly with GitHub (https://github.com), the online repository where we store the associated code and data. Because of the widespread use of CI in software development, alternative services that can run code on local or cloud-based computational infrastructure also exist (Beaulieu-Jones & Greene, 2017). We use CI to quality check data, test code using “unit tests” (Wilson et al., 2014), build models, make forecasts, and publicly present and archive the results (Figure 1b).

To ensure that software pipelines continue to run automatically as software dependencies change, a key component of “continuous analysis” is the use of a reproducible computational environment (Beaulieu-Jones & Greene, 2017). We followed Beaulieu and Greene’s (2017) recommendation to use software containers. Software containers are standalone packages that contain copies of everything needed to run a given piece of software, including the operating system (Boettiger, 2015). Once created, a software container is basically a time capsule, containing all the software dependencies in the exact state used to develop and run the software (Boettiger, 2015). If those dependencies change (or disappear) in the wider world, they still exist, unchanged, in the container. We use an existing platform, Docker (Merkel, 2014), to store an exact image of our complete software environment by adding our project specific code to a container created by the Rocker project, which is a Docker image with many important R packages (i.e., the tidyverse packages; Wickham, 2017) pre-installed (Boettiger & Eddelbuettel, 2017). We implemented this system because we experienced issues with external dependencies breaking our pipeline (e.g., when the `tscount` package (Liboschik et al., 2015), was temporarily removed from CRAN and could not be installed in the usual way). In combination, the automated running of the pipeline (continuous integration) and the guarantee it will not stop working unexpectedly due to software dependencies (via a software container) allows continuous analysis to serve as the glue that connects all stages of the forecasting pipeline.

## Data Collection, Entry, and Processing

Iterative forecasting benefits from frequently updated data so that state changes can be quickly incorporated into new forecasts (Dietze et al., 2018). Both frequent data collection and rapid processing are important for providing timely forecasts. Since we collect data monthly, ensuring that the models have access to the newest data requires a data latency period of less than 1 month from collection to availability for modeling. To accomplish this, we automated components of the data processing and quality assurance/quality control (QA/QC) process to reduce the time needed to add new data to the database [Yenni et al. (2018); Figure 1].

New data is double-entered into Microsoft Excel using the “data validation” feature. The two versions are then compared using an R script to control for errors in data entry. Quality control (QC) checks using the `testthat` R package (Wickham, 2011) are run on the data to test for validity and consistency both within the new data and between the new and archived data. The local use of the QC scripts to flag problematic data greatly reduces the time spent error-checking and ensures that the quality of data is consistent. The cleaned data is then uploaded to the GitHub-based PortalData repository (https://github.com/weecology/PortalData). GitHub (https://github.com/) is a software development tool for managing computer code development, but we have also found it useful for data management. On GitHub, changes to data can be tracked through the Git version control system which logs all changes made to any files in the repository, giving us a record of exactly of when specific lines of data were changed or added. All updates to data are processed through “pull requests,” which are notifications that someone has a modified version of the data to contribute. QA/QC checks are automatically run on the submitted data using continuous integration to ensure that no avoidable errors reach the official version of the dataset (Yenni et al., 2018).

We also automated the updating of supplementary data tables, including information on weather and trapping history, that were previously updated manually. As soon as new field data is merged into the repository, continuous integration updates all supplementary files. Weather data is automatically fetched from our cellular-connected weather station, cleaned, and appended to the weather data table. Supplementary data tables related to trapping history are updated based on the data added to the main data tables. Using CI for this ensures that all supplementary data tables are always up-to-date with the core data (Yenni et al., 2018).

## Data Sharing

The Portal Project has a long history of making its data publicly available so that anyone can use it for forecasting or other projects. Historically, the publication of the data was conducted through data papers (Ernest et al., 2009, 2016), the most common approach in ecology; this approach, however, caused years of data latency. With the recent switch to posting data directly to a public GitHub repository (Figure 1) with a CC0 waiver (i.e. no restrictions on data use; https://creativecommons.org/publicdomain/zero/1.0/), data latency for everyone has been reduced to less than one month, making meaningful iterative near-term forecasting possible for not only our group but other interested parties, as well (Ernest et al., 2018; Yenni et al., 2018).

## Data Manipulation

Once data is available, it must be processed into a form appropriate for modeling (Figure 1). For many ecological datasets, this requires not only simple data manipulation but also a good understanding of the data to facilitate appropriate aggregation. Data manipulation steps are often conducted using custom one-off code to convert the raw data into the desired form (Morris & White, 2013), but this approach has several limitations. First, each researcher must develop and maintain their own data manipulation code, which is inefficient and can result in different researchers producing different versions of the data for the same task. Subtle differences in data processing decisions have led to confusion when reproducing results for the Portal data in the past. Second, this kind of code is rarely robust to changes in data structure and location. Based on our experience developing and maintaining the Data Retriever (Morris & White, 2013; Senyondo et al., 2017), these kinds of changes are common. Finally, this kind of code is generally poorly tested, which can lead to errors based on mistakes in data manipulation. To avoid these issues for the Portal Project data, the Portal team has been developing an R package (portalr; http://github.com/weecology/portalr) for acquiring the data and handling common data cleaning and aggregation tasks. As a result, our modeling and forecasting code only needs to install this package and run the data manipulation and summary functions to get the appropriate data (Figure 1b). The package undergoes thorough automated unit testing to ensure that data manipulations are achieving the desired results. Having data manipulation code maintained in a separate package that focuses on consistently providing properly summarized forms of the most recent data has made maintaining the forecasting code itself much more straightforward.

## Modeling and Forecasting

Iterative near-term forecasting involves regularly updating a variety of different models (Figure 1). Ideally, new models should be easy to incorporate to allow for iterative improvements to the general modeling structure and approach. We use CI to update the models and make new forecasts each time the modeling code changes and when new data becomes available (Figure 1b). We use a modular plugin infrastructure to allow new models to be easily added to the system. This approach treats each model as an interchangable black box; all models have access to the same input data and generate the same structure for model outputs (Figure 2). Details of how to add a new model to the system are provided in the core GitHub repository (https://github.com/weecology/portalPredictions/wiki/Adding-a-new-model). During each run of the forecasting code, all existing models are run and the standardized outputs are combined into a single file to store the results of the different models’ forecasts. A weighted ensemble model is then added with weights based on how well individual models fit the training data. This plugin infrastructure makes it easy to add and compare very different types of models, from the basic time-series approaches currently implemented to the more complex state-space and machine learning models we hope to implement in the future. As long as a model script can load the provided data and produce the appropriate output, it will be run and its results incorporated into the rest of the forecasting system. This means that anyone can add a new model to the existing system by: 1) creating their own copy of the project (typically by forking the project on GitHub); 2) developing a new model; and 3) submitting a pull request to our repository.

**Figure 2:**
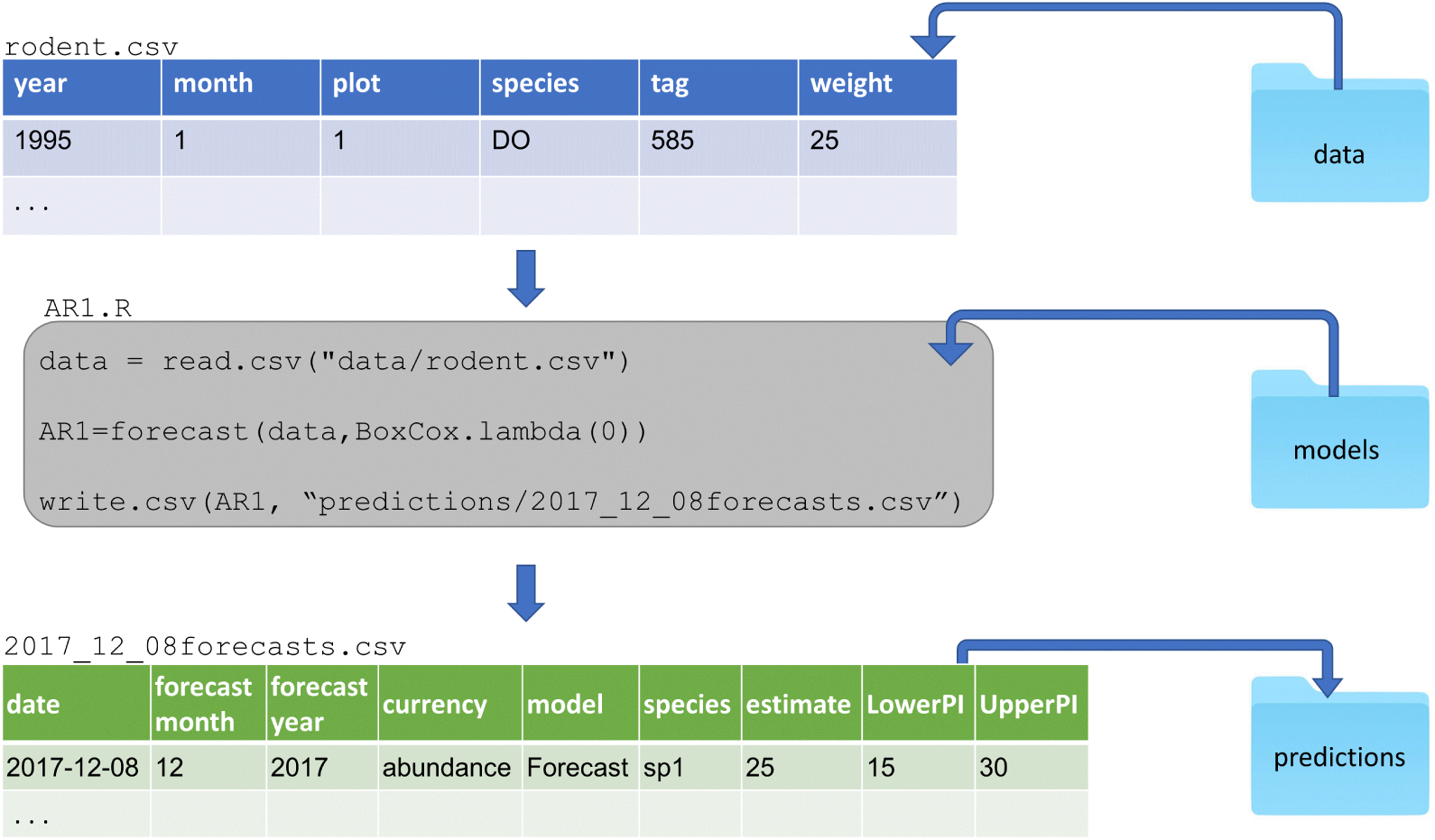
Demonstration of plugin infrastructure. All model scripts (represented here by the example AR1.R) are housed in a single folder. Each model script uses data provided by the core forecasting code (represented here by rodent.csv) and returns its forecast outputs in a predefined structure that is consistent across models (represented here by the example 2017_12_08forecasts.csv). Outputs from all models run on a particular date are combined into the same file (i.e. 2017_12_08forecasts.csv) to allow cross-model evaluations. Model output files are housed in a folder containing all forecast outputs from all previous dates to facilitate archiving and forecast assessment.

In addition to flexibility in what model structures can be supported, we also wanted to support flexibility in what the models predict. Allowing models to make forecasts for system properties ranging from individual species’ population abundances to total community biomass facilitates exploration of differences in forecastability across different aspects of ecological systems. We designed a forecast output format to support this. Each forecast output file contains the date being forecast, the collection date of the data used for fitting the models, the model name, the date the forecast was made, the state variable being forecast (e.g., rodent biomass, the abundance of a species), and the forecast value and associated uncertainty of that forecast (Figure 2). This allows us to store a variety of different forecasts in a common format and may serve as a useful starting point for developing a standard for storing ecological forecasts more generally.

Forecasts are currently evaluated using root mean square error (RMSE) to evaluate point forecasts and coverage to evaluate uncertainty. We plan to add additional metrics, like deviance, that incorporate both accuracy and uncertainty and better match the calibration method (Hooten & Hobbs, 2015; Dietze et al., 2018). In addition to evaluating the actual forecasts, we also use hindcasting (forecasting on already collected data; Jolliffe & Stephenson, 2003) to gain additional insight into the methods that work best for forecasting this system. For example, a model is fit using rodent observations up to June 2005, then used to make a forecast 12 months out to May 2006. The observations of that 12-month period can immediately be used to evaluate the model. Since hindcasting is conducted using data that has already been collected, it allows model comparisons to be conducted on large numbers of hindcasts and provides insight into which models make the best forecasts without needing to wait for new data to be collected (Harris et al., 2018). It can also be used to quickly evaluate new models instead of waiting for an adequate amount of data to accumulate. As the performance of different models is understood through evaluation of forecasts and hindcasts, models can be refined or removed from the system or ensemble to iteratively improve the resulting forecasts.

## Archiving

Publicly archiving forecasts before new data is collected allows the field to assess, compare, and build on forecasts made by different groups (McGill, 2012; Tredennick et al., 2016; Dietze et al., 2018; Harris et al., 2018) (Figure 1). Archiving serves as a form of pre-registration for model predictions because the forecasts cannot be modified once the data to assess them has been collected. This helps facilitate an unbiased interpretation of model performance. To serve this role, archives should be publicly accessible and be a permanent record that cannot be changed or deleted. This second criterion means that GitHub is not sufficient for archival purposes because repositories can be changed or deleted (Bergman, 2012; White, 2015). We explored three major repositories for archiving forecasts: FigShare (https://figshare.com/), Zenodo (https://zenodo.org/), and Open Science Framework (https://osf.io/). While all three repositories allowed for easy manual submissions (i.e., a human uploading files after each forecast), automating this process was substantially more difficult. Various combinations of repositories, APIs (i.e., interfaces for automatically interacting with the archiving websites), and associated R packages had issues with: 1) integrating authorization with continuous integration; 2) automatically making archived files public; 3) adding new files to an existing location; or 4) automatically permanently archiving the files. Our eventual solution was to leverage the GitHub-Zenodo integration (https://guides.github.com/activities/citable-code/) and automatically push forecasts to a GitHub repository from the CI server and release them via the GitHub API. The GitHub-Zenodo integration is designed to automatically create versioned archives of GitHub repositories. We created a repository for storing forecasts (https://github.com/weecology/forecasts) and linked this repository with Zenodo (a one-time manual process). Each time a new forecast is created, our pipeline adds the new forecasts to the GitHub repository and uses the GitHub API to create a new “release” for that repository. This triggers the GitHub-Zenodo integration, which automatically archives the resulting forecasts under a top-level DOI that refers to all archived forecasts (https://doi.org/10.5281/zenodo.839580). Through this process, we automatically archive every forecast made with a documented time-stamp. In addition, we also archive the full state of the modeling and forecasting repository (https://doi.org/10.5281/zenodo.833438). Through a similar process, the raw data in the data repository is also archived on a Zenodo whenever data is added or changed (Yenni et al., 2018), allowing retrieval of older versions of the data used for forecasting (https://doi.org/10.5281/zenodo.1219752). This ensures that every forecast is fully reproducible since the exact code and data used to generate every forecast is preserved. Early forecasts from this system are archived in the modeling and forecasting code archive, not in the newer repository ‘forecasts’.

## Presentation

Each month, we present our forecasts on a website that displays monthly rodent forecasts, model evaluation metrics, monthly reports, and information about the study site (Figure 3; http://portal.naturecast.org). The website includes a graphical presentation of the most recent month’s forecasts (including uncertainty) and compares the latest data to the previous forecasts. Information on the species and the field site are also included. The site is built using Rmarkdown (Allaire et al., 2017), which naturally integrates into the pipeline and is automatically updated after each forecast. The `knitr` R package (Xie, 2015) compiles the code into HTML, which is then published using Github Pages (https://pages.github.com/). The files for the website are stored in a subdirectory of the forecasting repository. As a result, the website is also archived automatically as part of archiving the forecast results.

**Figure 3:**
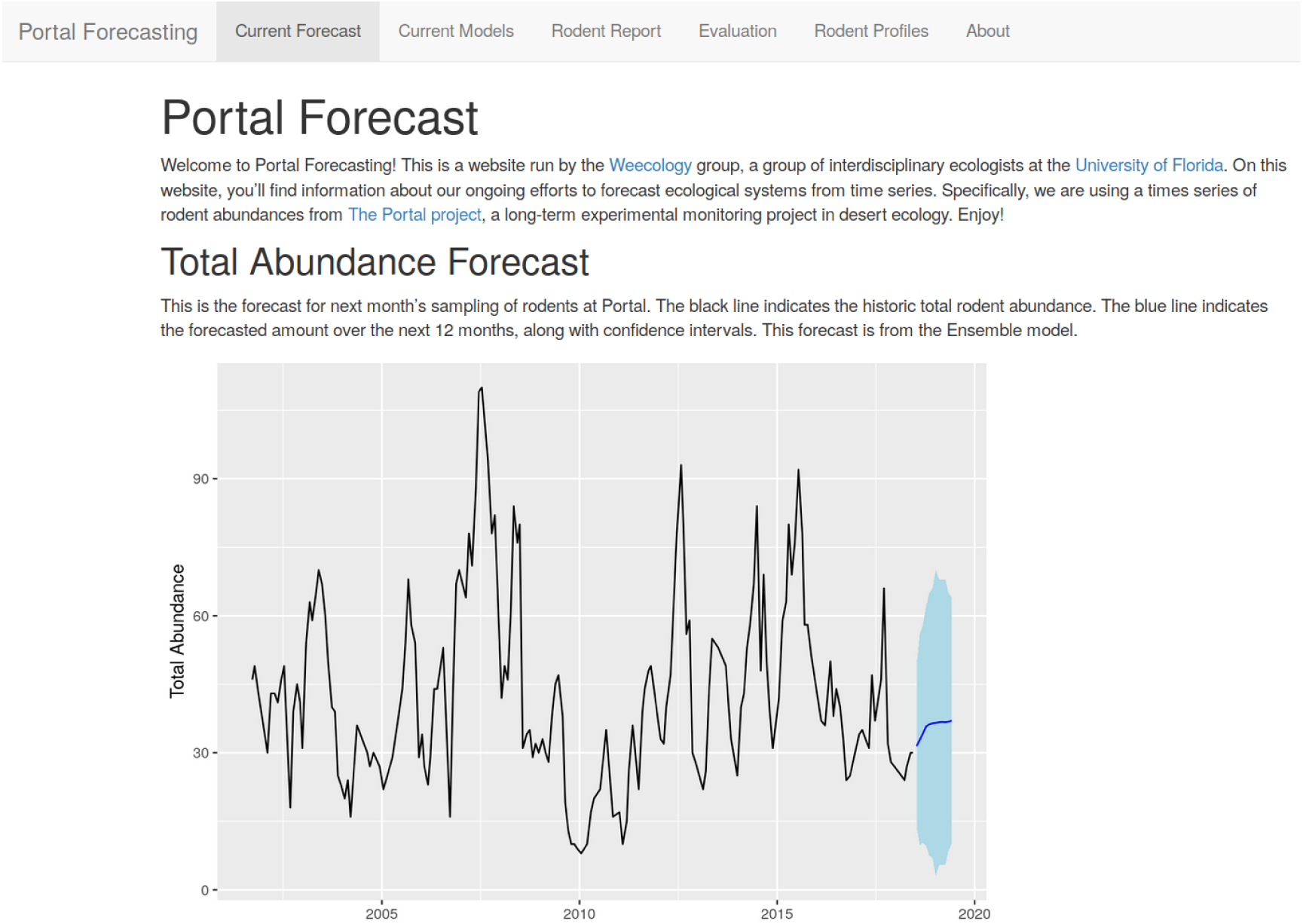
Screen capture of the homepage of the Portal Forecasting website (http://portal.naturecast.org). This site contains information on the most current forecasts, evaluation of forecast performance, and general information about the species being forecast.

## Discussion

Following the recommendations of Dietze et al (2018), we developed an automated iterative forecasting system (Figure 1) to support repeated forecasting of an ecological system. Our forecasting system automatically acquires and processes the newest data, updates the models, makes new forecasts, publicly archives those forecasts, and presents both the current forecast and information on how previous forecasts performed. Every week, the forecasting system generates a new set of forecasts with no human intervention, except for the entry of new field data. Our forecasting system ensures that forecasts based on the most recent data are always available and is designed to allow rapid assessment of the performance of multiple forecasting models for a number of different states of the system, including the abundances of individual species and community-level variables such as total abundance. To create this iterative near-term forecasting system, we used R to process data and conduct analyses and leveraged existing tools and services (i.e. GitHub, Travis, Docker) for more complicated cyberinfrastructure tasks. Thus, our approach to developing iterative near-term forecasting infrastructure provides an example for how short-term ecological forecasting systems can be developed.

We designed this forecasting system with the goal of making it relatively easy to build, maintain, and extend. We used existing technology for both running the pipeline and building individual components, which allowed us to build the system relatively cheaply in terms of both time and money. This included the use of tools like Docker for reproducibility, Travis CI continuous integration for automatically running the pipeline, Rmarkdown and `knitr` for generating the website, and the already existing integration between Github and Zenodo to archive the forecasts. By using this “continuous analysis” approach (Beaulieu-Jones & Greene, 2017), where analyses are automatically rerun when changes are made to data, models, or associated code, we have reduced the time required by scientists to run and maintain the forecasting pipeline. To make the system extensible so that new models could be easily incorporated, we used a plugin-based infrastructure so that adding a new model to the system is as easy as adding a single file to the ‘models’ folder in our repository (Figure 2). This should substantially lower the barriers to other scientists contributing models to this forecasting effort. We also automatically archive the resulting forecasts publicly so that the performance of these forecasts can be assessed by both us and other researchers as new data is collected. This serves as a form of pre-registration by providing a quantitative record of the forecast before the data being predicted was collected.

While building this system was facilitated by the use of existing technological solutions, there were still a number of challenges in making existing tools work for automated iterative forecasting. Continuous integration is designed primarily for running automated tests on software, not for running a coordinated forecasting pipeline. As a result, extra effort was sometimes necessary to figure out how to get these systems to work properly in non-standard situations, like running code that was not part of a software package. In addition, hosted continuous integration solutions, like Travis, provide only limited computational resources. As the number and complexity of the models we fit has grown, we have had to continually invest effort in reducing our total compute time so we can stay within these limits. Finally, we found no satisfactory existing solution for archiving our results. All approaches we tried had limitations when it came to automatically generating publicly-versioned archives of forecasts on a repeated basis, and our eventual solution was difficult to configure to such a degree that it will remain an impediment for most researchers. Overall, we found existing technology to be sufficient to the task of creating an iterative forecasting pipeline, but it required greater expertise and a greater investment of time than is ideal. Additional tool development to reduce the effort required for scientists to set up their own short-term forecasting systems would clearly be useful. Our efforts, however, show that it is possible to use existing tools to develop initial iterative systems as a method for both advancing scientific understanding and developing proof of concept forecasting systems.

Expanding the community of ecological forecasters using continous analysis approaches will require both an expansion of the current toolkit and the development of standards to facilitate interoperability of forecasts and forecasting systems. One of the major challenges for our current forecasting system is supporting computationally intensive forecasts. Projects involving larger datasets and/or complex modelling approaches will require either hosted solutions that provide infrastructure for running continuous integration and allow long-running distributed jobs or solutions that involve the user setting up their own continuous analysis sytem on cloud infrastructure or high performance computing centers. Event-driven serverless cloud platforms like OpenWhisk (https://openwhisk.apache.org/) and AWS Lambda (https://aws.amazon.com/lambda/) provide potential as hosted solutions, and open source continuous integrations systems like Jenkins (https://jenkins.io/) can be integrated with either cloud or high performance computing centers. However, both solutions are currently more complicated to set up than the hosted continuous integration approach we have employed using Travis. In addition to scalability issue for more computationally intensive projects, the toolkit for continuous analysis needs be made more researcher friendly. To broaden the user-base that can use continuous analysis for forecasting, we recommend the development of tools that make setting up continuous analysis easier by automating configuration steps. We also recommend the development of tools or data repository infrastructure to support the easy automated archiving of regularly generated data and forecasts (see Yenni et al., 2018). Finally, the development of standards for ecological forecasting to allow interoperability among forecasting systems will be essential for the growth of this field (see discussions of data standards, meta-data, and ontologies in ecology more broadly Jones et al., 2006; Madin et al., 2008; Michener & Jones, 2012). Now is an opportune time for developing these standards while the community of ecological forecasters is still small. While we have developed an initial format for storing and sharing forecasts, it is still lacking in several areas. Most notably, our approach to storing information on models and their associated uncertainty is insufficient for all but the simplest models. Improving this framework will require capturing both covariances between state variables and the full uncertainty in the models, by either storing full model objects or additional information like full ensembles of predictions (e.g., from Monte Carlo based approaches). This is challenging due to a lack of general standards for reporting uncertainty (Dietze et al., 2018).

Because of the breadth of expertise needed to set up our forecasting pipeline, our effort required a team with diverse skills and perspectives, ranging from software development to field site expertise. It is rare to find such breadth within a single individual, and our system was developed as a collaboration between the lab collecting and managing the data and a computational ecology lab. When teams have a breadth of expertise, communication can be challenging (Winowiecki et al., 2011). We found a shared base of knowledge related to both the field research and computational skills was important for the success of the group. The two labs are part of a joint interdisciplinary ecology group that has a mission of breaking down barriers between field and computational/theoretical ecologists (http://weecology.org). Everyone on the team had received training in fundamental data management and computing skills through a combination of university courses, Software and Data Carpentry workshops (Teal et al., 2015), and lab training efforts. In addition, everyone was broadly familiar with the study site and methods of data collection, and most team members had participated in field work at the site on multiple occasions. This provided a shared set of knowledge and vocabulary that actively facilitated interdisciplinary interactions. All members of the team actively participated in the development of the forecasting pipeline. Given the current state of tools for automated iterative forecasting, forecasting teams require some experience in working with continuous integration and APIs. This means either interdisciplinary teams or additional training will often be required for creating these pipelines until tool development improves. To improve the success of these diverse groups, we believe efforts at providing ‘team science’ training to scientists interested in forecasting will be beneficial for the success of iterative forecasting attempts for the foreseeable future (Read et al., 2016).

We developed infrastructure for automatically making iterative forecasts with the goals of making accurate forecasts for this well-studied system, learning what methods work well for ecological forecasting more generally, and improving our understanding of the processes driving ecological dynamics. The most obvious application of automated iterative ecological forecasting is for speeding up development of forecasting models by using the most recent data available and by quickly iterating to improve the models used for forecasting. By learning what works best for forecasting in this and other ecological systems, we will better understand what the best approaches are for ecological forecasting more generally. By designing the pipeline so that it can forecast many different aspects of the ecological community, we also hope to learn about what aspects of ecology are more forecastable. Finally, automated forecasting infrastructures like this one also provide a core foundation for faster scientific inquiry because new models can quickly be applied to data and compared to existing models. The forecasting infrastructure does the time-consuming work of data processing, data integration, and model assessment, allowing new research to focus on the models being developed and the inferences about the system that can be drawn from them (Dietze et al., 2018). We plan to use this pipeline to drive future research into understanding the processes that govern the dynamics of individual populations and the community as a whole. By regularly running different models for population and community dynamics, a near-term iterative pipeline such as ours should also make it possible to rapidly detect changes in how the system is operating, which should allow the rapid identification of ecological transitions or even possibly allow them to be prevented (Pace et al., 2017). By building an automated iterative near-term forecasting infrastructure, we can improve our ability to forecast natural systems, understand the biology driving ecological dynamics, and detect or even predict changes in system state that are important for conservation and management.

## Acknowledgements

We thank Henry Senyondo for help with continuous integration and Hao Ye for discussions and feedback on the manuscript. We thank all of the graduate students, postdocs, and volunteers who have collected the Portal Project over the last 40 years and the developers of the software and tools that made this project possible. We thank Heather Bradley for all of her logistical support that made this research possible. This research was supported by the National Science Foundation through grant 1622425 to S.K.M. Ernest and by the Gordon and Betty Moore Foundation’s Data-Driven Discovery Initiative through grant GBMF4563 to E.P. White.

## Data Accessibility

The data used in this study is from the Portal Project and is openly available (CC0) on GitHub (https://github.com/weecology/PortalData) and archived on Zenodo (Ernest et al. (n.d.)). Code for reproducing all analyses is available on GitHub (https://github.com/weecology/portalPredictions) and archived on Zenodo (White, Yenni, et al., 2018). Forecasts made by this system are all archived to Zenodo (White, Bledsoe, et al., 2018).

## Author Contributions

All authors conceived the ideas and designed methodology; All authors developed the automated forecasting system; EPW and SKME led the writing of the manuscript. All authors contributed critically to the drafts and gave final approval for publication.

## Box 1. Key practices for automated iterative near-term ecological forecasting

A list of some of the key practices developed by Dietze et al (2018) for facilitating iterative near-term ecological forecasting and discussion of why these practices are important.

### Data

#### 1. Frequent data collection

Frequent data collection allows models to be regularly updated and forecasts to be frequently evaluated (Dietze et al., 2018). Depending on the system being studied, this frequency could range from sub-daily to annual, but typically the more frequently the data is collected the better.

#### 2. Rapid data release under open licenses

Data should be released as quickly as possible (low latency) under open licenses so that forecasts can be made frequently and data can be accessed by a community of forecasters (Vargas et al., 2017; Dietze et al., 2018).

#### 3. Best practices in data structure

To reduce the time and effort needed to incorporate data into models, best practices in data structure should be employed for managing and storing collected data to ensure it is easy to integrate into other systems (interoperability) (Borer et al., 2009; Strasser et al., 2011; White et al., 2013).

### Models

#### 4. Focus on uncertainty

Understanding the uncertainty of forecasts is crucial to interpreting and understanding their utility. Models used for forecasting should be probabilistic to properly quantify uncertainty and to convey how this uncertainty increases through time. Evaluation of forecast models should include assessment of how accurately they quantify uncertainty as well as point estimates (Hooten & Hobbs, 2015; Harris et al., 2018).

#### 5. Compare forecasts to simple baselines

Understanding how much information is present in a forecast requires comparing its accuracy to simple baselines to see if the models yield improvements over the naive expectation that the system is static (Harris et al., 2018).

#### 6. Compare and combine multiple modeling approaches

To quickly learn about the best approaches to forecasting different aspects of ecology, multiple modeling approaches should be compared (Harris et al., 2018). Different modeling approaches should also be combined into ensemble models, which often outperform single models for prediction (Weigel et al., 2008).

## Cyberinfrastructure

In addition to improvements in data and models, iterative near-term forecasting requires improved infrastructure and approaches to support continuous model development and iterative forecasting (Dietze et al., 2018).

### 7. Best practices in software development

Best practices should be followed in the development of scientific software and modeling to make it easier to maintain, integrate into pipelines, and build on by other researchers. Key best practices include open licenses, good documentation, version control, and cross-platform support (Wilson et al., 2014; Hampton et al., 2015).

### 8. Support easy inclusion of new models

To facilitate the comparison and ensembling of different modeling approaches, code for fitting models and making forecasts should be easily extensible, to allow models developed by different groups to be integrated into a single framework (Dietze et al., 2018).

### 9. Automated end-to-end reproducibility

Each forecast iteration involves acquiring new data, updating the models, and making new forecasts. This should be done automatically without requiring human intervention. Therefore, the process of making forecasts should emphasize end-to-end reproducibility, including data, models, and evaluation (Stodden & Miguez, 2014), to allow the forecasts to be easily rerun as new data becomes available (Dietze et al., 2018).

### 10. Publicly archive forecasts

Forecasts should be openly archived to demonstrate that the forecasts were made without knowledge of the outcomes and to allow the community to assess and compare the performance of different forecasting approaches both now and in the future (McGill, 2012; Tredennick et al., 2016; Dietze et al., 2018; Harris et al., 2018). Ideally, the forecasts and evaluation of their performance should be automatically posted publicly in a manner that is understandable by both scientists and the broader stakeholder community.

## Box 2. Glossary of terms

**CI.** ‘Continuous Integration.’ The practice of continuouslly building and testing a code base as it is developed. **Data latency.** The time it takes for data to be available for use. **Docker.** An open-source Linux program for containerization (see software container). **git.** An open-source version control system. **GitHub.** A web-based host for git projects. Other options for a similar service include GitLab or Bitbucket. **PortalData.** The git repository for the Portal data, found on GitHub. **portalPredictions.** The git repository for the forecasts made using Portal data, found on GitHub. **portalr.** An R package for using the Portal data. **QA/QC.** ‘Quality Assurance.’ Testing the quality of a product. ‘Quality Control.’ The process of ensuring the quality of a product. **Rocker.** A project making it easy to use Docker containers in the R environment. **Software container.** Allows a developer to package up an application with all of the parts it needs to run reliably. **testthat.** R package used to set up automated testing for QA/QC. **Travis.** A continuous integration service that integrates easily with GitHub and R. Examples of similar programs are Jenkins or CodeShip. **Unit test.** A component of quality control in which each smallest testable part of software is formally tested. **Zenodo.** An open data archive that integrates easily with GitHub.

